# The origins and molecular evolution of sperm

**DOI:** 10.1101/2025.11.03.685759

**Authors:** Arthur Matte, David Skerrett-Byrne, Andrea Manica, Adria C. LeBoeuf

**Author notes:** **Author correspondence** AM, ACL.

## Abstract

Sperm is a nearly universal cell type among animals, yet its evolutionary origins remain unclear. Using comparative proteomics and phylogenomics across 32 species, we reconstruct the 700+ million year evolutionary history of sperm to define the Last Universal Common Sperm (LUCS), a conserved core of 301 gene families enriched in motility and energy metabolism. We found that most of the LUCS toolkit was developed in filozoan unicellular ancestors approximately 400 Ma before animals, revealing sperm as a reconfigured legacy from our unicellular past rather than an invention of multicellularity. Sperm shows a within-cell evolutionary gradient, both spatial and temporal, where ancient proteins dominate the tail, while younger innovations concentrate in the head. The oldest sperm components are disproportionately associated with human infertility, establishing an empirical bridge between evolutionary depth and clinical relevance. Together, our findings reveal how evolutionary history is inscribed within a single cell and can guide clinical insights.

## Introduction

Gametes are the cellular basis of sexual reproduction in animals. While eggs contain genetic material and the resources for early development, sperm face a fundamentally different challenge: they must transport genetic material out of one body, locate an egg, and deliver their cargo through fusion to create a zygote. Across Metazoa, sperm have undergone extreme morphological and molecular evolution, shaped by fertilization environments and sexual selection ^1–3^. The cellular mechanics of sperm are increasingly well described ^4–9^, yet a fundamental question remains unresolved: what are the evolutionary origins of the sperm cell itself?

Do sperm share a conserved molecular toolkit? When was this toolkit first assembled, and how has it been remodeled over ∼700 million years of animal evolution? Do different sperm compartments (*e*.*g*., motile tails, egg-binding heads) follow distinct evolutionary trajectories? And despite sperm’s fundamental role, functions still break down in the form of infertility ^10,11^. How does this fragility intersect with the evolutionary history of sperm?

To address these questions, we integrated comparative proteomics across 32 animal species with phylogenomics spanning 62 opisthokonts, tracing sperm evolution in its genomic and proteomic context. This approach allowed us to reconstruct the ancestral sperm toolkit, resolve the diversification of this cell type across lineages and compartments, and test whether its most ancient components disproportionately underlie human infertility. Our results reveal sperm as a reconfigured unicellular legacy with a spatial and temporal evolutionary gradient within a single cell, and they establish the Last Universal Common Sperm (LUCS) as both a molecular and clinical reference for understanding sperm evolution and infertility.

## Results

### Sperm molecular composition across animals

To investigate the molecular basis of sperm evolution, we conducted a large-scale comparative proteomic analysis across 32 metazoan species, capturing over 700 million years of sperm evolution. Using species-specific genome assemblies for peptide-spectrum matching, we identified a total of 59,710 proteins, an average of 1,865 ± 269 per species. We clustered these proteins into 7,864 orthogroups, on a backbone of orthology built from 62 genomes spanning non-opisthokont eukaryotes to metazoans. These orthogroups represent gene families defined by sequence homology and inferred evolutionary relationships.

Comparison of orthogroup composition across species revealed distinct layers of conservation and divergence across sperm proteomes (Fig. 1). At the broadest level, a deeply conserved core of orthogroups makes up sperm of nearly all species. Surrounding this universal set are clade-specific ‘sub-cores’ marking major evolutionary branches. Beyond these, species-specific innovations have emerged, alongside a more diffuse layer of orthogroups distributed sporadically across lineages without clear phylogenetic clustering. This stratification suggests that sperm are built on shared indispensable molecular architecture and have sustained lineage-specific diversifications.

**Figure 1.**
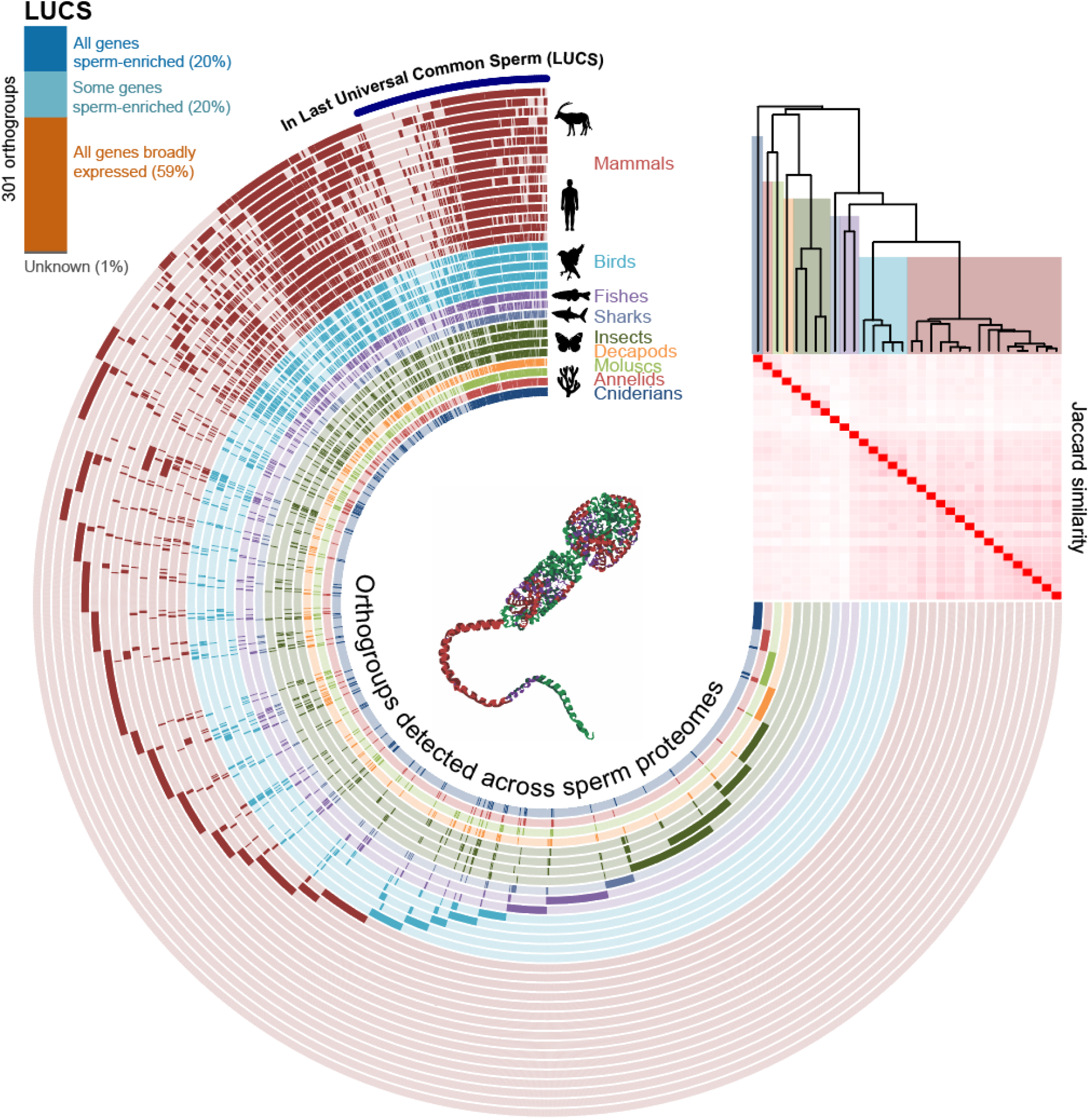
Comparative landscape of sperm proteomes across Metazoa. Presence-absence matrix of orthogroups detected across sperm proteomes from 32 species. Each column represents a species and each row an orthogroup. Only the 500 most abundant orthogroups per species are shown to account for variability in proteome completeness. The 301 orthogroups in LUCS (Last Universal Common Sperm) are highlighted by the purple bar around the matrix exterior and details of their specificity to sperm is given by the bar in the top-left corner. Orthogroup expression was classified according to transcriptomic data from the Human Protein Atlas, where combined expression in testis and epididymis was compared against other tissues to identify orthogroups composed exclusively of sperm-enriched genes, broadly expressed genes or a mixture of both.

### Tracing where sperm began: The Last Universal Common Sperm (LUCS)

To investigate the molecular origins of sperm, we used parsimony to reconstruct orthogroup recruitment and loss across sperm proteomes of 32 species, identifying 634 orthogroups predicted to be present in sperm at the metazoan root. To ensure robustness of our inferences, we retained only orthogroups consistently predicted to be present in sperm at the root in ≥95% of 1,000 jackknife replicates (each excluding 10% of species). We define this conservative consensus of 301 orthogroups as the Last Universal Common Sperm (LUCS) (Fig. 1).

To determine whether these gene families are merely fundamental cellular machinery as opposed to true signatures of sperm, we analysed the expression patterns of the human genes present in the LUCS orthogroups and performed functional enrichment against the background of all gene symbols detected across all species’ sperm (Fig. S1). Across the LUCS genes, 59% are broadly expressed across human tissues and are enriched in core metabolic functions such as mitochondrial respiration. In contrast, 40% of LUCS genes have sperm-biased expression (fully or partially sperm-enriched expression) and are enriched in cell motility functions such as cilium movement and axonemal structure (Fig. S1). The remaining 1% of orthogroups are absent in human sperm, lack annotation, or lack HPA data, with no information available about their expression pattern.

### Sperm-like cells before animals and the pre-metazoan origin of the sperm toolkit

When and how did the LUCS molecular toolkit emerge? While the genomic foundations of animal multicellularity are well studied ^e.g., 12–15^, the evolutionary origin of gametes remains unresolved. Did the evolution of sperm involve the invention of novel cellular machinery, or instead the co-option of pre-existing machinery?

Tracing orthogroup origins across eukaryotes showed that only ∼1% of LUCS orthogroups are metazoan-specific. The vast majority of LUCS orthogroups predate animals, supporting earlier work indicating that sperm draw primarily on ancient cellular machinery ^16,17^. However, this observation alone does not reveal when these genes first underwent coordinated innovation.

To pinpoint timing, we reconstructed the innovation histories of the 301 LUCS orthogroups across the opisthokont phylogeny using a parsimony-based framework. Expansions at each ancestral node were then benchmarked against a background of 30,000 randomly chosen orthogroups, allowing us to distinguish genuine bursts from noise. If sperm was a metazoan invention, we would expect a burst of innovation in these LUCS gene families at the metazoan root ^18^. Instead, the innovation burst in LUCS orthogroups occurred much earlier, in the last common ancestor of Filozoa, the clade containing Metazoa, Choanoflagellata, and Filasterea (Fig. 2, Fig. S2).

**Figure 2.**
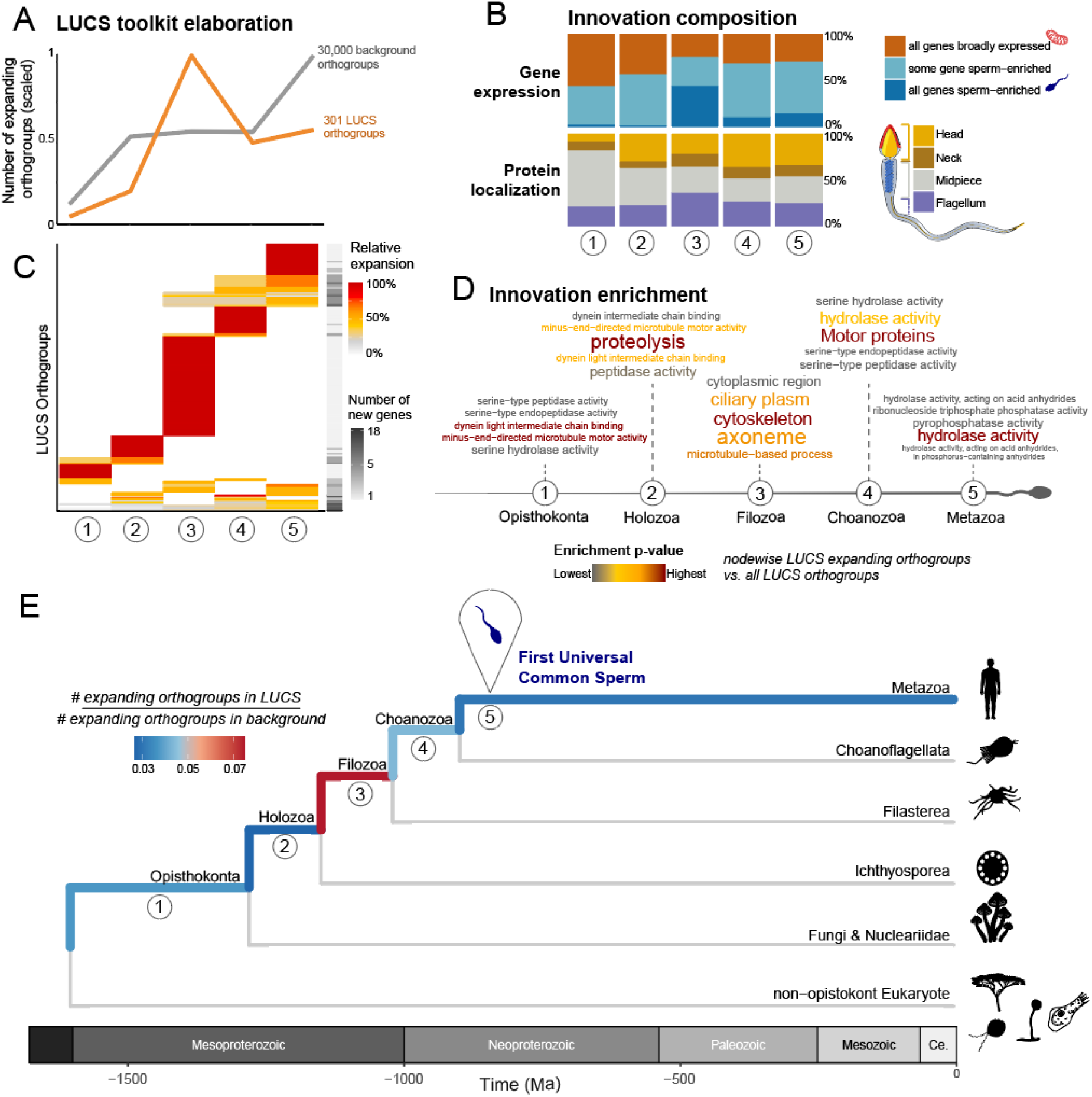
Pre-metazoan assembly of the sperm toolkit. Nodewise analysis across the opisthokont phylogeny shows a major expansion of LUCS orthogroups in a filozoan ancestor. (**A**) Number of expanding LUCS orthogroups compared to a random background of 30,000 orthogroups. (**B)** Details of protein expression patterns and subcellular localizations of expanding orthogroups at each node. (**C**) Relative magnitude of expansions of each LUCS orthogroup per node. (**D**) Nodewise functional enrichment of expanding orthogroups. Word lists indicate the most enriched term from innovating orthogroups vs. all LUCS orthogroups. (**E**) Evolutionary pathway of sperm from the root of Opisthokonta to Metazoa, with branch color denoting the number of expanding orthogroups in LUCS compared to a random background of 30,000 orthogroups.

Functional enrichment of expanding orthogroups at each node relative to all LUCS orthogroups, showed that the Filozoa innovation burst was characterized by flagellar architecture (*e*.*g*., cytoskeleton, axoneme), while subsequent innovations at the Choanoflagellata node, then Metazoa, were primarily related to enzymatic activities (*e*.*g*., hydrolase, serine peptidases) (Fig. 2C, Fig. S3). This points to a scenario in which an ancient swimming, flagellated unicellular blueprint was established in a Filozoa ancestor and was eventually co-opted into sperm.

The pre-metazoan deployment of this fundamental sperm toolkit is evident in extant filozoans: choanoflagellates use this toolkit for swimming, feeding, and even anisogamous gamete fusion ^15,19,20^. Together, these findings reveal that the fundamental architecture of sperm was largely assembled before the origin of animals and later co-opted into reproduction, while other cells diversified into novel, non-motile somatic forms.

Our inference of a pre-metazoan origin of the sperm toolkit is necessarily conservative for several reasons. The lack of available sperm proteomes from sponges and ctenophores reduces resolution at the base of Metazoa. This 120 Ma gap pushes LUCS toolkit age estimations toward younger origins by including proteins recruited in the Planulozoa, yet our results indicate that the strongest burst of innovation in LUCS is far older. Similarly, parsimony-based ancestral reconstruction, though robust to uneven sampling and long divergence times, underestimates gene duplication and loss. This acts as a buffer rather than an artifact: it compresses complexity but does not generate spurious bursts of innovation. The strong expansion we detect at the filozoan root is therefore unlikely to be an artifact of our methods.

### A within-cell evolutionary gradient: a tail of conservation and a head of innovation

Building on the LUCS framework, we wanted to understand how sperm proteomes diversified across animals. While sperm and testis are known hotspots of rapid protein evolution ^21,22^, and subcellular compartments have been found to experience distinct evolutionary pressures ^5,23,24^, a phylogenetically resolved breakdown of protein ages across sperm has so far been lacking.

We used parsimony-based reconstructions to trace the recruitment age of each orthogroup into sperm proteomes. Across 32 species, an average of 38 ± 0.1% of sperm proteins traced back to LUCS, while 15 ± 0.1% were species-specific (Fig. 3A) with stable proportions across taxa despite variation in proteomic sampling depth (Fig. S4). The weak phylogenetic signals for both the LUCS and species-specific fractions (Pagel’s λ of 0.65 and 0.19, respectively) suggest a broadly conserved pattern: an ancient foundational frame of conserved metabolic and motility machinery, layered by lineage-specific innovations. Importantly, the genomic origin of a gene did not predict proteomic recruitment. Some orthogroups that arose >1.5 billion years ago entered sperm only recently (*e*.*g*., ITPR1^25^), while some younger families were recruited as soon as they were found in the genome (*e*.*g*., DEFB22^26,27^) (Fig. 3B), indicating that genomic age alone can misrepresent cell-type evolution.

**Figure 3.**
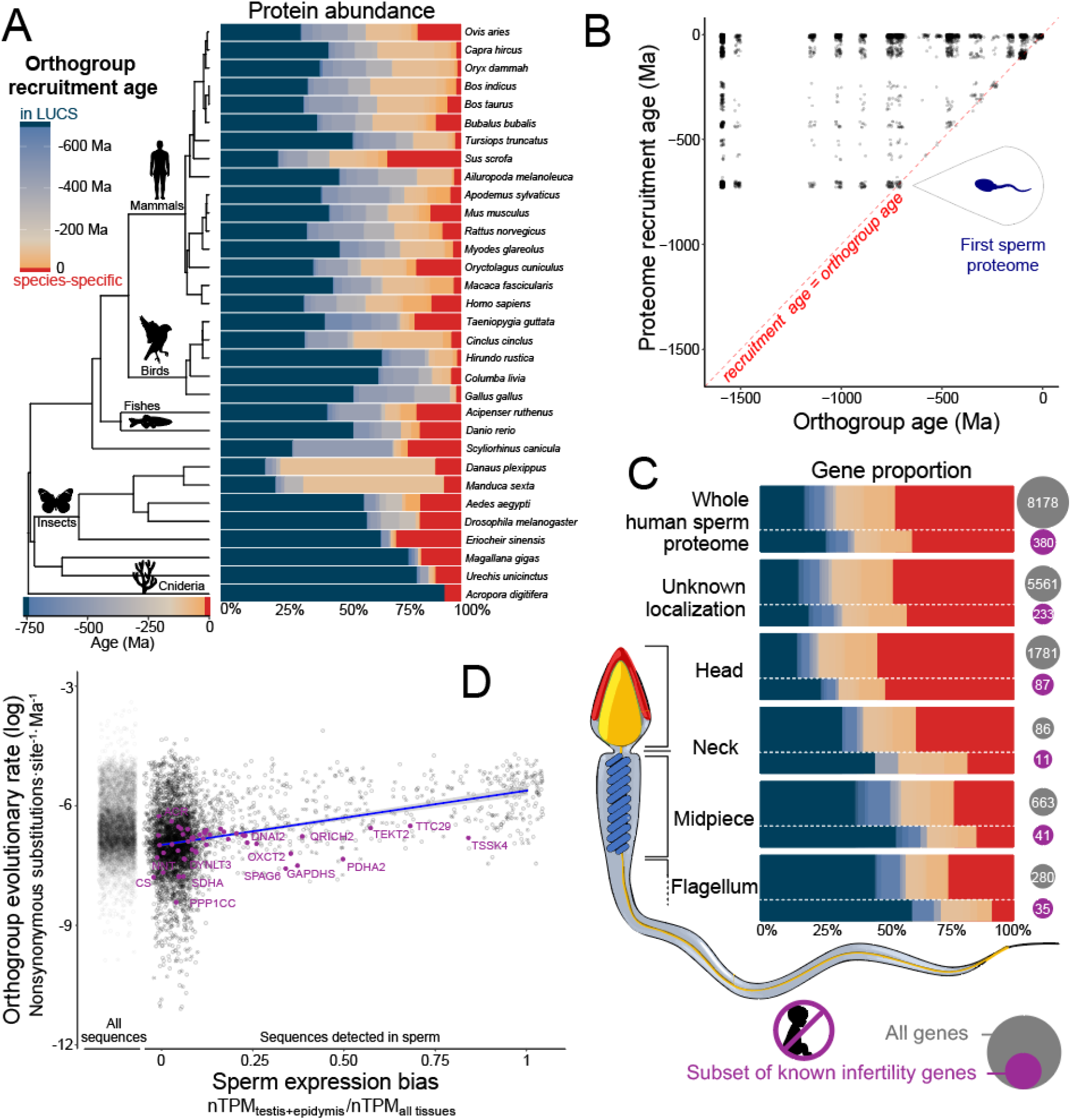
Sperm proteomes integrate ancient foundations with rapid, lineage-specific innovations. **(A)** Relative abundance of proteins of different evolutionary ages across species. Note that the age estimate of specific-specific genes is constrained by the species set. **(B)** Comparison of genomic origin vs. proteomic recruitment age in the sperm proteomes. Dotted red lines indicate when genomic age equals proteomic age (*i*.*e*., an orthogroup emerged in the genome and was immediately recruited into sperm). **(C)** Gradient of protein recruitment age along the sperm head-to-tail axis. Subcompartment highlights the age proportion for the subset of genes known for monogenic infertility, revealing a systematic bias toward older origins. Genes present in multiple compartments (*e*.*g*., head and neck) were represented multiple times in the localization breakdown. **(D)** Orthogroup evolutionary rate and sperm-biased expression. Orthogroup evolutionary rates were estimated as the median number of non-synonymous substitutions per site per million years (visualised as log-transformed), from pairwise sequence comparisons across species using only sequences detected in sperm. Sperm-biased expression corresponds to the average ratio nTPM_testis+epididymis_/nTPM_across all tissues_, from the Human Protein Atlas database. The left dot cloud of “All sequences” corresponds to evolutionary rates calculated for all orthogroups present in humans using all sequences present across all species for comparison. Purple dots and gene names highlight genes known for causing monogenic infertility belonging to orthogroups in LUCS.

We next tested whether evolutionary age varies along the sperm axis. Using Human Protein Atlas annotations to assign sperm proteins to subcellular compartments revealed a clear head-to-tail gradient. The tail retains ancient, conserved modules (*e*.*g*., DNAH10^28^), while the head is dominated by recently recruited, lineage-specific proteins (*e*.*g*., IZUMO3 ^29^) (linear model, *p* < 0.001, R^2^= 0.10, Fig. 3C). This gradient aligns with functional constraints. The tail acts as a near-universal engine for motility, while the head, and in particular the acrosome, engages directly with the egg surface proteins in a molecular arms race driven by sexual selection and reproductive isolation ^24^.

This molecular gradient contrasts with sperm morphology, where head length tends to evolve slowly relative to other compartments ^30^, likely under stabilizing structural constraints (*e*.*g*., hydrodynamism). Our findings add nuance to this picture: while the physical architecture of the sperm head remains conserved, its molecular composition is at the cutting edge of evolutionary change.

Altogether, these findings establish sperm as a modular system, with a conserved flagellar core and an innovation-prone head. By resolving both temporal and spatial stratification, our phylogenetic framework shows how evolutionary history is differentially inscribed within a single cell.

### Sperm-specific orthogroups diverge faster than non-specific ones

Reproductive proteins are known to evolve rapidly ^5,21,23^. How this pattern integrates with deep evolutionary time has yet to be explored. To address this, we compared orthogroup sperm-expression bias and evolutionary rate. For each orthogroup, sperm-expression bias was defined as the ratio of normalized transcript abundance (nTPM) in testis and epididymis relative to total expression across 50 tissues. Evolutionary rates were quantified as the average number of nonsynonymous substitutions per site per million years, calculated from pairwise comparisons of all sperm-detected sequences across the 32 species.

Although broadly expressed orthogroups display a wide range of evolutionary rates, orthogroups with stronger testis and epididymis specificity evolved markedly faster (*linear model, p* < 0.001, R^2^= 0.11, Fig. 3D). Orthogroups exclusively expressed in sperm were predicted to evolve 3.9-fold faster (95% CI: 3.5-4.3) than non-sperm-biased orthogroups, consistent with previous work ^21,31^.

Together, these results demonstrate that rapid divergence in sperm proteins stems primarily from regulatory restriction: when expression is confined to sperm, genes escape pleiotropic constraint and can become hotspots of adaptive innovation.

### Ancient sperm genes and infertility

Given this stratification, we asked whether genes that have been conserved for longer evolutionary periods are more likely to be essential for sperm function, as reflected by their association with infertility. We compared the age distribution of 380 sperm-localized genes linked to monogenic male infertility ^32^ to that of the full sperm proteome.

The age distribution of infertility genes differed significantly from background expectation (χ^2^= 21.34, df = 2, *p* < 0.001) with a systematic bias toward old origins; this was true proteome-wide, but also within every sperm compartment, with flagellum and neck harboring the highest proportion of infertility genes (12.5% and 12.8% respectively) (Fig. 3C). Infertility loci were 1.5-fold enriched among the most ancient LUCS orthogroups and primarily under-represented among human-specific genes (152 vs. 177 expected, 0.86-fold downregulation). These results establish an empirical bridge between evolutionary depth and clinical relevance. The disruption of genes conserved since LUCS likely compromises the core machinery shared by all animal sperm, whereas lineage-specific proteins may provide adaptation to reproductive contexts whose disruption is less likely to cause complete infertility.

To facilitate clinical and evolutionary exploration, we provide an annotated sperm proteome for the 32 species studied (Table S2) and humans (Table S3), linking each detected protein to its evolutionary stratum, subcellular localization, expression profile, and infertility association. This resource highlights conserved, unexplored candidates with high potential relevance to male reproductive health.

## Discussion

By providing a phylogenetically resolved reconstruction of sperm evolution, our results reveal sperm as a repurposed pre-metazoan unicellular body plan rather than a product of multicellularity. Whether this evolutionary continuity is unique to sperm or shared with other cell types remains unknown. Other animal cell types may also trace back to ancient modules or represent true novelties of multicellularity, a distinction that only future comparative work can resolve.

Sperm’s unicellular origin aligns with proposals that the germline preserves ancestral unicellular features ^33,34^. The pre-metazoan deployment of the sperm blueprint also raises further questions: is animal sperm homologous to choanoflagellate gametes ^15^, a surviving specialization of a choanoflagellate-like swimming state, or a co-option of shared machinery into a new reproductive context? Such questions intersect with the deeper challenge that cell types are evolutionary continua, not discrete boxes ^35,36^. Nonetheless, theoretical models predicting that multicellularity emerged to sustain and protect more specialized reproductive cells ^37,38^ resonate strongly with this view. From this perspective, soma can be considered a cooperative scaffold evolved to sustain and transmit fundamentally unicellular entities.

Our results reveal a striking within-cell evolutionary gradient. Ancient components dominate the tail, powering low-Reynolds-number motility, whereas the head accumulates recent innovations at the fertilization interface. This spatial and temporal layering of conservation and novelty exemplifies a broader rule of biological organization: indispensable cores endure, while interfaces diversify. The same logic governs neurons, which conserve synaptic scaffolds but continually reinvent receptors ^39^ and immune cells, which preserve signaling cascades but diversify recognition machinery ^40^. Sperm, by making this duality visible along a single cellular axis, provides a powerful model and illustrates how evolutionary history is spatially inscribed within cells.

This framework carries immediate biomedical implications. The oldest sperm proteins are disproportionately linked to human infertility, establishing an empirical bridge between evolutionary depth and clinical relevance. If evolutionary depth predicts indispensability in sperm, this may extend to other cell types such as neurons, immune cells, and beyond. Evolutionary reconstruction thus not only illuminates cell-type history but also identifies prime molecular targets for clinical intervention.

## Materials and Methods

### Proteomic data sourcing

Proteomic datasets of mature spermatozoa were retrieved from the ProteomeXchange Consortium (www.proteomexchange.org), including data from PRIDE, iProX, and MassIVE repositories ^41–44^. We excluded samples containing immature sperm or seminal fluids, and for experiments with treatment groups, retained only control or wild-type samples. We also excluded low-confidence datasets following Pini et al. ^9^ (*i*.*e*., with aberrant patterns and too low protein discovery). This curation yielded 54 datasets representing 32 species and ∼2 TB of spectral data. Proteomic dataset sources and associated metadata are provided in Supplementary Table S1.

### Genomic data sourcing and orthology inferences

Genomic sequences were obtained primarily from NCBI RefSeq to ensure consistent annotation. Species were selected to match available sperm proteomes, with additional representatives from Porifera, Ctenophora, and Cnidaria included to improve resolution at deep nodes. For *Urechis unicinctus* (Echiura, Annelida), we used the published author-annotated genome ^45^. To extend analyses beyond Metazoa, we incorporated non-metazoan opisthokonts and other eukaryotic outgroups, supplementing the limited Filasterea representation with three annotated genomes from Ocaña-Pallarès et al. ^46^. Gene orthology was inferred from these 62 genomes using OrthoFinder v2.5.4 ^47,48^ using the longest isoform per gene as representative following guidance from OrthoFinder. We verified that non-RefSeq genomes did not yield outlier results (Fig. S4-S5). Genome source details and metadata are provided in Supplementary Table S1.

### Proteome processing

Raw spectral data were processed in MaxQuant (version 2.0.1.0)^49^ using species-specific genome annotations. Search analyses were performed per species and study, with default settings except for setting a minimum of one unique peptide per protein, up to three missed cleavages (Trypsin/P), and enabling “match between runs.” Default false discovery rate (FDR) thresholds of 1% were applied at both the peptide and protein levels. Protein abundance was estimated using iBAQ (intensity-based absolute quantification) ^50^ values, calculated as the sum of all peptide intensities divided by the number of theoretically observable peptides per protein. Downstream filtering excluded known contaminants, reverse hits, zero-abundance proteins, and proteins detected in only one sample when replicates were available. iBAQ values were normalized within samples, then averaged per species and normalized within species. Resulting matched sequences and annotated sperm proteomes are provided in Table S2 (all 32 species) and Table S3 (human focus).

### Time-calibrated species phylogeny

A dated species tree was obtained from the TimeTree of Life database (https://timetree.org/), which integrates divergence estimates from published studies ^51^.

To reflect current phylogenetic understanding ^52^, we repositioned Ichthyosporea as diverging before Filasterea and incorporated relationships among Filasterea species from Ocaña-Pallarès et al. ^46^, as these were not resolved in TimeTree. These manual adjustments do not affect downstream ancestral state reconstructions, which depend solely on tree topology and are insensitive to branch length.

### Ancestral proteome and genome reconstruction

We modeled the evolutionary history of all orthogroups detected across sperm proteomes using maximum parsimony implemented in *asr_max_parsimony (*castor package v1.8.3) ^53^. For each orthogroup, we inferred ancestral presence or absence at all internal nodes of the phylogeny. Because parsimony can yield multiple equally optimal solutions, we applied a custom cost matrix that weighted gene gains 1.1× higher than losses. This mild asymmetry resolved ties by slightly favoring loss, consistent with general expectations of gene evolution, without imposing a strong directional bias ^54^.

Orthogroup recruitment age was defined as the earliest inferred presence in sperm along the tree. In parallel, we reconstructed orthogroup birth and death across genomes using the same method to compare genomic age with recruitment into the sperm proteome (Fig. 3B).

Maximum parsimony was preferred over likelihood-based methods because it is non-model-based, avoids reliance on uncertain branch lengths, and limits equivocal reconstructions at deep nodes over long evolutionary timescales ^55^.

To assess robustness in age estimations, we performed jackknife resampling with 1,000 replicates, each randomly excluding 10% of species. Age estimates were highly consistent, with low coefficients of variation (Fig. S6A). Deviations were mainly confined to loosely represented orthogroups, consistent with recent acquisitions: omitting particular species occasionally shifted assignments toward species-specific or slightly older ages (Fig. S6A).

Classical phylostratigraphic methods ^56,57^ assign gene ages based on the deepest taxonomic level at which homologs can be detected, often using BLAST or HMMER similarity searches. For our purposes, genomic age alone does not necessarily reflect when a gene was recruited into a particular cell type (Fig 3B). Phylostratigraphy cannot capture this distinction, whereas our orthogroup-based ancestral state framework reconstructs both gene birth and recruitment into the sperm proteome. This approach is deliberately more conservative, but it ensures that inferred evolutionary strata reflect cell-type histories rather than genomic origin only, providing a more accurate foundation for reconstructing sperm evolution.

### LUCS inference

To define the Last Universal Common Sperm (LUCS), we built upon our ancestral proteome reconstruction analysis with a deliberately conservative strategy. While 665 orthogroups were initially inferred at the metazoan root, we restricted LUCS to the 301 consistently recovered in >95% of 1,000 jackknife replicates (Fig. S6B), thereby minimizing artifacts from species representation. A broader threshold (≥85% replicates) expanded LUCS to 634 orthogroups, but all core evolutionary and functional signals presented in this study were fully captured by the 301-stringent set. This principled filtering ensured that LUCS reflects a robust ancestral core rather than an artifact of taxon sampling, positioning it as a stable molecular reference point for sperm evolution.

To further resolve when LUCS orthogroups assembled during eukaryotic evolution, we reconstructed gene copy number histories across the Opisthokonta phylogeny using maximum parsimony implemented in *asr_max_parsimony* (*Casor* package v1.8.3) ^53^.

Transition costs were set to “proportional”, permitting all state changes but increasing costs with the magnitude of the change.

At each ancestral node, we counted orthogroups with at least one inferred duplication. These values were then standardized against a null distribution of 30,000 randomly sampled orthogroups using the function *sample* from R ^58^, yielding a normalized measure of sperm toolkit expansion across evolutionary time.

### Gene ontology, enrichment analysis, and infertility linkage

Gene symbols corresponding to primary isoforms matched with sperm proteomes were retrieved from NCBI annotations. Functional enrichment analyses were conducted using the gprofiler2 package (version 0.2.3) ^59^, using false discovery rate (FDR) correction for multiple testing ^60^.

Infertility-associated genes were obtained from Houston et al. ^32^, the most comprehensive curated database of monogenic causes of male infertility to our knowledge. Genes detected in the human sperm proteome were grouped into three evolutionary strata (in LUCS, intermediate age, or human-specific) based on their inferred orthogroup recruitment age (see above). For enrichment testing, the background was defined as all genes detected in the human sperm proteome, and the foreground as the subset of these genes associated with infertility. To assess whether infertility-associated genes were non-randomly distributed across evolutionary strata, we compared the observed counts of infertility-associated genes with those expected based on the overall distribution of all detected genes using a chi-square goodness-of-fit test.

### Gene expression patterns

Tissue-specific expression of human sperm proteins was assessed using the Consensus RNA dataset from the Human Protein Atlas (HPA) (version 24.0; https://www.proteinatlas.org), which integrates transcriptomic data from HPA, GTEx, and FANTOM5 across 50 tissues ^61^.

We adapted the HPA classification framework to assess sperm specificity by combining expression in testis and epididymis. For each gene, expression across these tissues was summed and compared against other tissues, assigning genes into one of two categories: 1) Sperm-enriched: Combined expression in testis and epididymis ≥ 4× higher than the average expression across all other tissues; 2) Broadly expressed: Genes not meeting the above criteria.

To evaluate expression patterns at the orthogroup level, we summarized classifications across all genes in each orthogroup. Orthogroups were categorized as: 1) All genes are sperm-enriched; 2) All genes are broadly expressed; 3) Some genes are sperm-enriched, some are broadly expressed. No expression information could be obtained for genes lacking NCBI annotation and/or HPA transcriptomic data, and were left as unknown.

We additionally calculated orthogroup sperm expression bias as the average ratio nTPM_testis+epididymis_ / nTPM_across all tissues_, from all genes of a given orthogroup.

This simplified framework enabled a systematic assessment of how specifically sperm proteome genes, and their orthogroups, are expressed in male reproductive tissues. Gene- and orthogroup-level classifications for the human sperm proteome are reported in Table S2.

### Sperm evolution across subcellular compartments

Sperm compartments fulfill distinct roles and may evolve under different selective pressures. To test for heterogeneity in recruitment age, we assigned proteins to anatomical compartments along the sperm longitudinal axis.

For each primary isoform detected in the human sperm proteome, we searched its gene symbol in the Human Protein Atlas database (https://www.proteinatlas.org/) and retrieved its “*Main subcellular locations*” ^62^. Annotations flagged as “uncertain” were excluded to focus on “approved” and “supported” annotations. After this filtering, we analyzed 2631 out of 8210 sperm proteins. Localizations were grouped into four major compartments: 1) Head: acrosome, equatorial segment, perinuclear theca, calyx, nucleus; 2) Neck / Connecting piece: connecting piece, basal body; 3) Midpiece: mitochondria, midpiece, annulus (junction between midpiece and principal piece); 4) Flagellum: principal piece, axoneme, flagellum, flagellum centriole, end piece. To assess longitudinal gradients, we coded compartments ordinally (1–4) and fit a linear model with recruitment age as the response variable and compartment position as the predictor.

Gene compartments and subcellular localization for the human sperm proteome are available in Table S2. Several orthogroups (9.1%) contained genes co-occurring in several compartments (Fig. S7). For figure and statistical analysis, these orthogroups were duplicated to be present in every category independently.

### Estimating sperm protein evolutionary rate

Protein evolutionary rates were estimated from within-orthogroup pairwise amino-acid sequence comparisons. Sequences identified in sperm proteomes were aligned in R using *pairwiseAlignment* (pwalign package, v1.2.0) ^63^ with the BLOSUM62 substitution matrix. Alignment columns containing gaps in either sequence were removed, and the proportion of mismatched residues over ungapped sites was taken as the substitution fraction. To avoid unreliable estimates, alignments with more than 45% gapped positions were excluded. Species separation times were obtained from the study phylogeny and used to normalize sequence divergence, yielding substitution rates per site per million years. This approach is more appropriate for large scale comparisons because it quantifies sequence turnover directly, independent of selection models such as dN/dS ^64^.

Sequence substitution rates were averaged within orthogroups. For orthogroups containing multiple paralogs (and hence multiple species pairwise comparisons), we used the mean of the minimum (most conservative) and maximum (least conservative) pairwise distances across species pairs. Orthogroup mean evolutionary rates were log-transformed and regressed against orthogroup mean sperm expression bias (see above) using a linear model. Data distributions were compared with background evolutionary rates calculated from all orthogroups across the 62 genomes analyzed in this study (Fig. 3C).

## Acknowledgment

The authors gratefully acknowledge all authors of the datasets utilised, without whom this work would not be possible. We thank David MacKenzie for his spermatozoa illustration. This work was supported by the Balfour PhD Studentship to A. Matte, a HFSP grant to ACL (RGP0023/2022), a Swiss National Science Foundation grant (PR00P3_179776) to ACL, and a National Health and Medical ResearchCouncil of Australia (NHMRC) Emerging Leadership Fellowship (APP2034392) to DSB.

## Supplementary materials

**Table S1**. Details of publicly available proteomic and genomic datasets investigated in this study.

**Table S2**. Annotated mature sperm proteome for the 32 species investigated in this study, with details of matched sequences, associated gene symbol, iBAQ, predicted orthogroup recruitment age in the sperm proteome, and predicted orthogroup birth age along the Opisthokonta phylogeny.

**Table S3**. Subset of Table S2 focusing on human sperm proteome, with additional details on gene expression, subcellular localization, and link with infertility.

## Notes

### Competing Interest Statement

The authors have declared no competing interest.

